## Supplementary Material for "Decoding subjective emotional arousal from EEG during an immersive Virtual Reality experience"

### Supplementary Methods

#### Details of the rollercoasters

The "Space" rollercoaster did not feature outstanding events during the ride besides two vertical spins starting around 47 s and 73 s after the onset of the experience. Virtual collisions of asteroids floating through the scenery led to explosions of the celestial bodies involved, accompanied by an explosive sound. Apart from this, there were little sound effects during the space experience.

The "Andes" rollercoaster included a steep drop (24 s after onset), two jumps with steep landings (31 s and 67 s after onset), two passages through fires under the tracks (20 and 55 s after onset) and a looping (60 s after onset). Sound effects mimicked the sound of the waggon on the tracks, the fire, and the airflow. In the background a jingling melody was played.

#### Simulator sickness questions

The wording and items to assess simulator sickness, were:

Please rate on a scale from 1 to 7 how much each symptom below is affecting you right now:

(A) General discomfort

(B) Nausea

(C) Dizziness

(D) Headache

(E) Blurred vision

(F) Difficulty concentrating

### Supplementary Figures

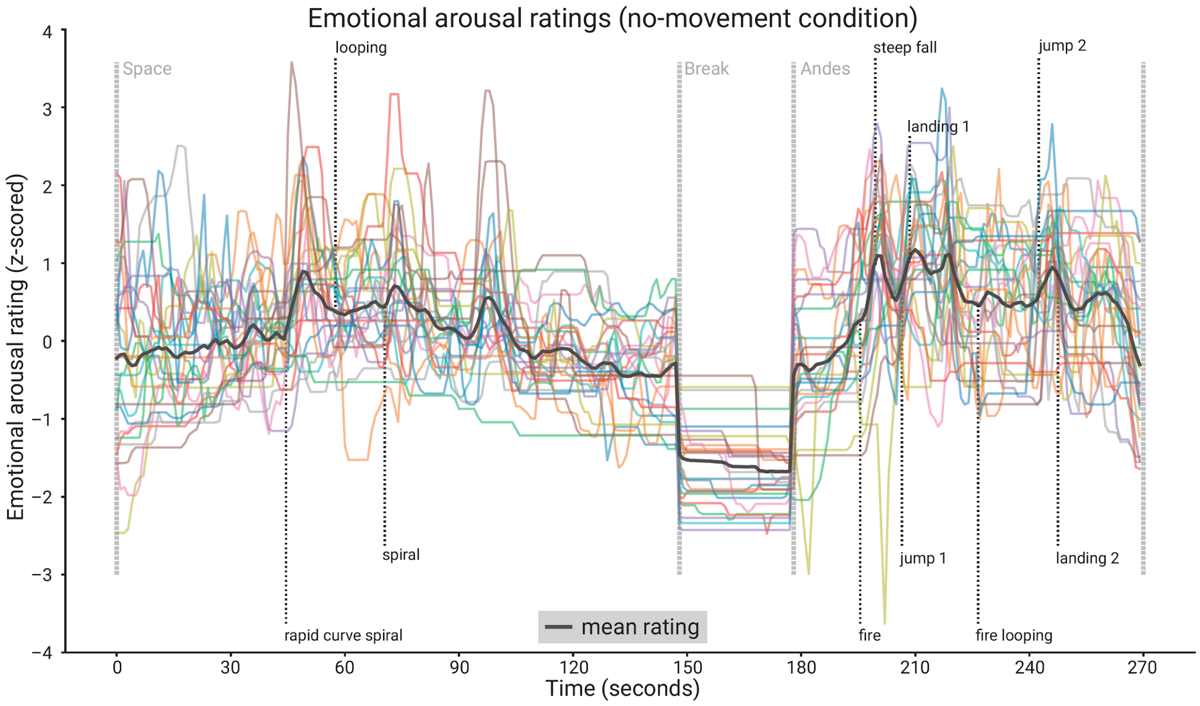


**Figure 5 – figure supplement 1. Subjective emotional arousal ratings (no-movement condition).** Emotional arousal ratings of the experience (without head movement). Coloured lines: individual participants; black line: mean across participants; vertical lines (light grey): beginning of the three phases (Space Coaster, Break, Andes Coaster); vertical lines (dark grey): manually labelled salient events (for illustration).

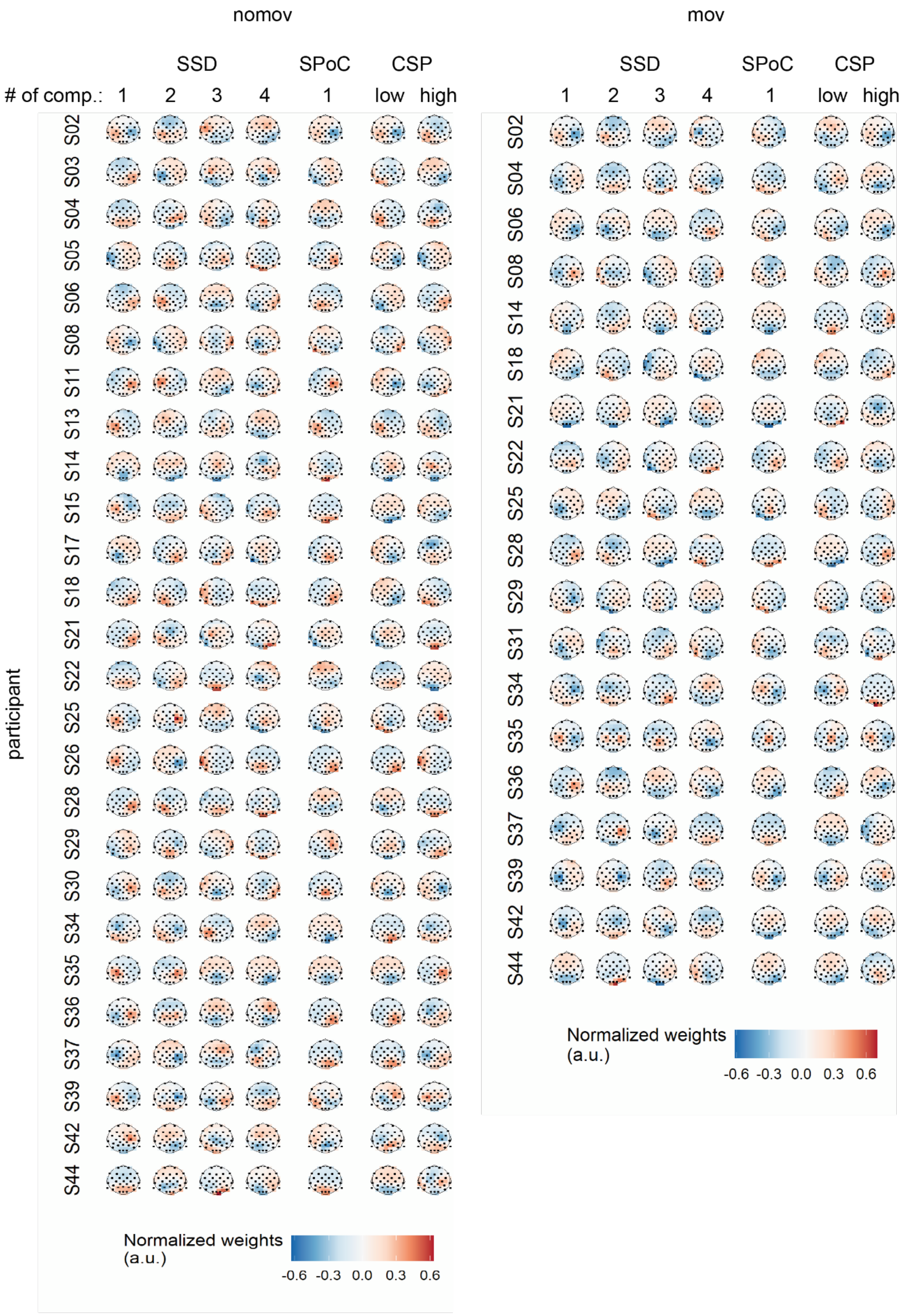
nomov
mov
SSD
SPoC
CSP
### of comp.: 1 2 3 4 1 low high
participant
S02 S03 S04 S05 S06 S08 S11 S13 S14 S15 S17 S18 S21 S22 S25 S26 S28 S29 S30 S34 S35 S36 S37 S39 S42 S44
S02 S04 S06 S08 S14 S18 S21 S22 S25 S28 S29 S31 S34 S35 S36 S37 S39 S42 S44
Normalized weights (a.u.)
-0.6 -0.3 0.0 0.3 0.6

4

**Figure 6 – figure supplement 1. Spatial patterns per single subject and movement condition yielded by the different spatial signal decompositions.** For SSD the four patterns corresponding to the four highest eigenvalues among the accepted components (see methods) are displayed (note: subjects with *less* than four accepted SSD components were discarded for further analysis; for subjects with *more* than four accepted components, all of these components went into the further analyses but only the first four patterns are shown here). For SPoC the pattern associated with the component that yielded the strongest correlation between target and source power is displayed. For CSP the patterns associated with the components that maximized power during states of low and high emotional arousal are shown.

### Supplementary Tables

| Subject | $SPoC_\lambda$ | $SPoC_r$ | $SPoC_p$ | $CSP_{acc}$ | $CSP_p$ | $LSTM_{acc}$ | $LSTM_p$ |
|---|---|---|---|---|---|---|---|
| S02 | 0.081 | 0.099 | 0.999 | 0.683 | 0.000 | 0.683 | 0.000 |
| S03 | -0.194 | -0.224 | 0.401 | 0.617 | 0.002 | 0.633 | 0.000 |
| S04 | -0.405 | -0.245 | 0.040 | 0.617 | 0.002 | 0.561 | 0.117 |
| S05 | -0.934 | -0.180 | 0.127 | 0.672 | 0.000 | 0.600 | 0.009 |
| S06 | -0.266 | -0.281 | 0.032 | 0.639 | 0.000 | 0.572 | 0.062 |
| S08 | -0.185 | -0.239 | 0.155 | 0.517 | 0.709 | 0.583 | 0.030 |
| S11 | -0.408 | -0.219 | 0.022 | 0.594 | 0.014 | 0.611 | 0.004 |
| S13 | -0.328 | -0.218 | 0.386 | 0.539 | 0.333 | 0.533 | 0.412 |
| S14 | -0.210 | -0.332 | 0.013 | 0.650 | 0.000 | 0.572 | 0.062 |
| S15 | -0.227 | -0.288 | 0.086 | 0.550 | 0.205 | 0.583 | 0.030 |
| S17 | -1.088 | -0.336 | 0.018 | 0.567 | 0.086 | 0.594 | 0.014 |
| S18 | -0.254 | -0.201 | 0.098 | 0.533 | 0.412 | 0.533 | 0.412 |
| S21 | -0.322 | -0.274 | 0.337 | 0.722 | 0.000 | 0.672 | 0.000 |
| S22 | -0.639 | -0.204 | 0.053 | 0.572 | 0.062 | 0.522 | 0.602 |
| S25 | -1.491 | -0.531 | 0.000 | 0.600 | 0.009 | 0.589 | 0.021 |
| S26 | -0.464 | -0.276 | 0.303 | 0.778 | 0.000 | 0.656 | 0.000 |
| S28 | -0.215 | -0.153 | 0.128 | 0.617 | 0.002 | 0.556 | 0.157 |
| S29 | -0.360 | -0.170 | 0.148 | 0.472 | 0.502 | 0.589 | 0.021 |
| S30 | -0.432 | -0.198 | 0.737 | 0.672 | 0.000 | 0.644 | 0.000 |
| S34 | -0.813 | -0.527 | 0.000 | 0.589 | 0.021 | 0.617 | 0.002 |
| S35 | -0.442 | -0.259 | 0.166 | 0.694 | 0.000 | 0.650 | 0.000 |
| S36 | -0.771 | -0.411 | 0.012 | 0.656 | 0.000 | 0.644 | 0.000 |
| S37 | -0.665 | -0.335 | 0.095 | 0.650 | 0.000 | 0.561 | 0.117 |
| S39 | -0.147 | -0.185 | 0.732 | 0.489 | 0.823 | 0.539 | 0.333 |
| S42 | -0.357 | -0.261 | 0.219 | 0.517 | 0.709 | 0.539 | 0.333 |
| S44 | -0.492 | -0.231 | 0.172 | 0.611 | 0.004 | 0.611 | 0.004 |

**Figure 10 – figure supplement 1. Results per decoding approach and participant (nomov condition).** *SPoC:* values for the component with the smallest (i.e., most negative) correlation between its alpha power and the emotional arousal ratings. *λ:* covariance; *r*: Spearman correlation coefficient; *p*: *p*-values obtained from the permutation test (see *Methods*) on the single subject level; *CSP* and *LSTM*: classification results. *acc*: proportion of correctly classified samples across the cross-validation folds. *p*: *p*-values obtained from the exact binomial test on the single subject level (see *Methods*).

| Subject | $SPoC_\lambda$ | $SPoC_r$ | $SPoC_p$ | $CSP_{acc}$ | $CSP_p$ | $LSTM_{acc}$ | $LSTM_p$ |
|---|---|---|---|---|---|---|---|
| S02 | 0.029 | 0.042 | 0.978 | 0.556 | 0.157 | 0.550 | 0.205 |
| S04 | -0.228 | -0.188 | 0.262 | 0.578 | 0.044 | 0.594 | 0.014 |
| S06 | -0.155 | -0.201 | 0.302 | 0.556 | 0.157 | 0.594 | 0.014 |
| S08 | -0.354 | -0.249 | 0.212 | 0.656 | 0.000 | 0.622 | 0.001 |
| S14 | -0.132 | -0.225 | 0.098 | 0.533 | 0.412 | 0.589 | 0.021 |
| S18 | -0.358 | -0.264 | 0.063 | 0.567 | 0.086 | 0.617 | 0.002 |
| S21 | -0.493 | -0.157 | 0.462 | 0.561 | 0.117 | 0.589 | 0.021 |
| S22 | -0.567 | -0.369 | 0.000 | 0.483 | 0.709 | 0.544 | 0.263 |
| S25 | -1.496 | -0.527 | 0.001 | 0.711 | 0.000 | 0.711 | 0.000 |
| S28 | -0.473 | -0.217 | 0.039 | 0.556 | 0.157 | 0.583 | 0.030 |
| S29 | -0.591 | -0.319 | 0.006 | 0.578 | 0.044 | 0.583 | 0.030 |
| S31 | -0.342 | -0.302 | 0.006 | 0.650 | 0.000 | 0.539 | 0.333 |
| S34 | -0.051 | -0.077 | 0.925 | 0.717 | 0.000 | 0.667 | 0.000 |
| S35 | -0.177 | -0.272 | 0.087 | 0.578 | 0.044 | 0.622 | 0.001 |
| S36 | -0.375 | -0.271 | 0.095 | 0.689 | 0.000 | 0.633 | 0.000 |
| S37 | -1.117 | -0.464 | 0.029 | 0.600 | 0.009 | 0.661 | 0.000 |
| S39 | -0.364 | -0.374 | 0.004 | 0.667 | 0.000 | 0.656 | 0.000 |
| S42 | -0.108 | -0.197 | 0.625 | 0.633 | 0.000 | 0.633 | 0.000 |
| S44 | -0.567 | -0.264 | 0.083 | 0.678 | 0.000 | 0.656 | 0.000 |

**Figure 10 – figure supplement 2. Results per decoding approach and participant (mov condition).** Variables like in Figure 10–figure supplement 1.

| Subject | resting state | nomov | mov |
|---|---|---|---|
| S01 | 11.322 | NaN | NaN |
| S02 | 10.715 | 10.787 | 11.190 |
| S03 | 10.535 | 10.711 | NaN |
| S04 | 10.097 | 9.865 | 10.241 |
| S05 | 10.299 | 9.969 | 10.236 |
| S06 | 10.277 | 12.293 | 12.503 |
| S07 | 11.589 | 10.701 | NaN |
| S08 | 11.194 | 11.450 | 11.056 |
| S09 | 9.961 | 10.917 | 11.905 |
| S10 | 9.423 | NaN | NaN |
| S11 | 11.396 | 11.831 | 9.349 |
| S13 | 11.185 | 9.633 | 10.535 |
| S14 | 10.482 | 9.300 | 10.573 |
| S15 | 10.271 | 11.498 | NaN |
| S16 | 10.554 | NaN | NaN |
| S17 | 9.711 | 9.080 | 11.424 |
| S18 | 8.830 | 10.740 | 10.384 |
| S19 | 11.082 | NaN | NaN |
| S20 | 10.355 | 11.131 | 12.139 |
| S21 | 10.717 | 12.405 | 12.307 |
| S22 | 9.403 | 10.823 | 10.307 |
| S23 | 9.413 | NaN | NaN |
| S24 | 10.876 | 9.701 | 8.825 |
| S25 | 11.034 | 11.489 | 11.283 |
| S26 | 10.993 | 11.146 | NaN |
| S27 | 10.475 | 11.686 | 12.253 |
| S28 | 9.118 | 11.026 | 11.161 |
| S29 | 9.975 | 11.586 | 11.893 |
| S30 | 9.247 | 10.195 | NaN |
| S31 | 9.011 | 10.050 | 11.070 |
| S32 | 9.733 | NaN | NaN |
| S33 | 8.758 | NaN | NaN |
| S34 | 10.879 | 12.205 | 12.246 |
| S35 | 9.689 | 10.087 | 10.568 |
| S36 | 10.067 | 10.254 | 10.316 |
| S37 | 9.499 | 10.588 | 9.942 |
| S38 | 8.570 | NaN | NaN |
| S39 | 10.065 | 10.961 | 11.096 |
| S40 | 9.924 | NaN | NaN |
| S41 | 9.712 | NaN | NaN |
| S42 | 10.393 | 11.840 | 12.111 |
| S43 | 10.121 | NaN | NaN |
| S44 | 10.564 | 10.376 | 10.853 |
| S45 | 10.324 | NaN | NaN |

**Figure 2 – figure supplement 1. Selected alpha peaks (8-13 Hz) per participant and condition.**

Results of FOOF for three conditions: eyes-closed resting state, nomov, and mov.

| Subject | LSTM | FC | l.rate | reg. | reg. strength | activ.func. | components |
|---|---|---|---|---|---|---|---|
| S02 | 30,20 | 0 | 5e-4 | l2 | 1.44 | elu | 1,2 |
| S03 | 20,10 | 0 | 1e-3 | l1 | 0.72 | elu | 1,2 |
| S04 | 30,30 | 30 | 5e-4 | l1 | 1.44 | relu | 1,2,3,4,5,6,7,8 |
| S05 | 50,40 | 10 | 5e-4 | l2 | 0.00 | relu | 1,2,3,4 |
| S06 | 30,25 | 0 | 5e-4 | l1 | 0.36 | elu | 1,2,3,4,5,6,7 |
| S08 | 80,50 | 25 | 1e-3 | l1 | 0.00 | relu | 1,2,3 |
| S11 | 30,25 | 0 | 5e-4 | l1 | 0.36 | elu | 1,2,3,4,5,6,7 |
| S13 | 40,20 | 10 | 5e-4 | l1 | 0.00 | relu | 1 |
| S14 | 30,25 | 0 | 5e-4 | l1 | 0.36 | elu | 1,2,3,4,5,6,7 |
| S15 | 40,15 | 0 | 1e-3 | l1 | 0.72 | relu | 1,2,3,4,5 |
| S17 | 30,30 | 30 | 5e-4 | l1 | 1.44 | relu | 1,2,3,4,5,6,7,8 |
| S18 | 30,25 | 0 | 5e-4 | l1 | 0.36 | elu | 1,2,3,4,5,6,7 |
| S21 | 30,25 | 0 | 5e-4 | l1 | 0.36 | elu | 1,2,3,4,5,6,7 |
| S22 | 30,25 | 0 | 5e-4 | l1 | 0.36 | elu | 1,2,3,4,5,6,7 |
| S25 | 20,15 | 10 | 1e-3 | l1 | 0.00 | relu | 1,2,3,4,5,6,7 |
| S26 | 40,15 | 0 | 1e-3 | l1 | 0.72 | relu | 1,2,3,4,5 |
| S28 | 30,20 | 0 | 5e-4 | l2 | 1.44 | elu | 1,2 |
| S29 | 50,40 | 10 | 5e-4 | l2 | 0.00 | relu | 1,2,3,4 |
| S30 | 30,30 | 30 | 5e-4 | l1 | 1.44 | relu | 1,2,3,4,5,6,7,8 |
| S34 | 30,25 | 0 | 5e-4 | l1 | 0.36 | elu | 1,2,3,4,5,6,7 |
| S35 | 100100 | 0 | 5e-4 | l1 | 0.00 | elu | 1,2,3,4 |
| S36 | 20,15 | 10 | 1e-3 | l1 | 0.00 | relu | 1,2,3,4,5,6,7 |
| S37 | 30,25 | 0 | 5e-4 | l1 | 0.36 | elu | 1,2,3,4,5,6,7 |
| S39 | 40,20 | 10 | 5e-4 | l1 | 0.00 | relu | 1 |
| S42 | 30,20 | 0 | 5e-4 | l2 | 1.44 | elu | 1,2 |
| S44 | 40,30 | 10 | 5e-4 | l2 | 0.72 | elu | 1,2,3,4,5,6,7 |

**Figure 4 – figure supplement 1. LSTM hyperparameter search (nomov condition).** *LSTM*: number of cells per layer. *FC*: number of hidden units in fully connected layer, before final output neuron. *l.rate*: learning rate. *reg.*: type of weight regularizer. *reg. strength*: respective regularization strength. *activ.func*: intermediate layer activation function. *components*: individually selected components for training after SSD selection. *mean accuracy*: mean classification accuracy across 10 folds of cross validation.

| Subject | LSTM | FC | l.rate | reg. | reg. strength | activ.func. | components |
|---|---|---|---|---|---|---|---|
| S02 | 65,30 | 25 | 5e-4 | l2 | 0.18 | relu | 1,2,3,4,5,6,8 |
| S04 | 40,15 | 10 | 1e-3 | l2 | 1.44 | relu | 1,2,3,4,6 |
| S06 | 30,25 | 0 | 1e-3 | l2 | 0.00 | elu | 1,2,3,4,5,6,7 |
| S08 | 65,30 | 25 | 5e-4 | l2 | 0.18 | relu | 1,2,3,4,5,6,7 |
| S14 | 50,50 | 15 | 1e-2 | l1 | 0.00 | elu | 1,2,3,4,5,6,7,8 |
| S18 | 15,15 | 10 | 1e-3 | l2 | 0.72 | relu | 1 |
| S21 | 40,15 | 10 | 1e-3 | l2 | 1.44 | relu | 1,2,3,4,6 |
| S22 | 80 | 0 | 5e-4 | l2 | 0.18 | elu | 1,2 |
| S25 | 15,15 | 10 | 1e-3 | l2 | 0.72 | relu | 1 |
| S28 | 30,25 | 0 | 1e-3 | l2 | 0.00 | elu | 1,2,3,4,5,6,7 |
| S29 | 65,30 | 25 | 5e-4 | l2 | 0.18 | relu | 1,2,3,4,5 |
| S31 | 40 | 0 | 1e-2 | l2 | 0.18 | elu | 1,2,3,4 |
| S34 | 65,30 | 25 | 5e-4 | l2 | 0.18 | relu | 1,2,3,4,5 |
| S35 | 65,30 | 25 | 5e-4 | l2 | 0.18 | relu | 1,2,3,4,5,6,8 |
| S36 | 65,30 | 25 | 5e-4 | l2 | 0.18 | relu | 1,2,3,4,5,6,8 |
| S37 | 65,30 | 25 | 5e-4 | l2 | 0.18 | relu | 1,2,3,4,5,6,8 |
| S39 | 65,30 | 25 | 5e-4 | l2 | 0.18 | relu | 1,2,3,4,5 |
| S42 | 10 | 0 | 1e-3 | l2 | 0.72 | relu | 1 |
| S44 | 65,30 | 25 | 5e-4 | l2 | 0.18 | relu | 1,2,3,4,5 |

**Figure 4 – figure supplement 2. LSTM hyperparameter search (mov condition).** Variables like in Figure 4–figure supplement 1.
